# Torques within and outside the human spindle balance twist at anaphase

**DOI:** 10.1101/2023.12.10.570990

**Authors:** Lila Neahring, Yifei He, Nathan H. Cho, Gaoxiang Liu, Jonathan Fernandes, Caleb J. Rux, Konstantinos Nakos, Radhika Subramanian, Srigokul Upadhyayula, Ahmet Yildiz, Sophie Dumont

## Abstract

At each cell division, nanometer-scale motors and microtubules give rise to the micron-scale spindle. Many mitotic motors step helically around microtubules in vitro, and most are predicted to twist the spindle in a left-handed direction. However, the human spindle exhibits only slight global twist, raising the question of how these molecular torques are balanced. Here, using lattice light sheet microscopy, we find that anaphase spindles in the epithelial cell line MCF10A have a high baseline twist, and we identify factors that both increase and decrease this twist. The midzone motors KIF4A and MKLP1 are redundantly required for left-handed twist at anaphase, and we show that KIF4A generates left-handed torque in vitro. The actin cytoskeleton also contributes to left-handed twist, but dynein and its cortical recruitment factor LGN counteract it. Together, our work demonstrates that force generators regulate twist in opposite directions from both within and outside the spindle, preventing strong spindle twist during chromosome segregation.

## Introduction

At each cell division, the micron-scale spindle self-organizes from nanometer-scale molecular components to divide the genome. While the identities of nearly all these building blocks are known (Neumann et al., 2010), many questions remain about how they together give rise to the architecture, mechanics, and function of the spindle as an ensemble. Mitotic motors, over a dozen species of which are present in the human spindle, illustrate this gap: although the motility and force-generating capacity of many motors have been closely studied in vitro (Canty et al., 2021; Cross and McAinsh, 2014), it remains poorly understood how motors cooperate in the dense microtubule network of the spindle to give rise to larger-scale microtubule architecture.

Several motors have been found to generate net torque on microtubules in vitro, resulting in helical motility around the microtubule track. Mitotic motors have rotational pitches ranging from approximately 0.3-2.3 µm (Walker et al., 1990; Yajima et al., 2008), far more extreme than the supertwist of microtubules (no supertwist in 13-protofilament microtubules and a slight left-handed twist with ∼6 µm pitch in 14-protofilament microtubules) (Ray et al., 1993). The torques produced by kinesins that crosslink and slide microtubule pairs are sufficiently strong to twist and coil two microtubules around each other (Mitra et al., 2020). The plus-end-directed yeast kinesin-8 Kip3 (Bormuth et al., 2012; Mitra et al., 2018) and *Caenorhabditis elegans* kinesin-6 ZEN-4 (Maruyama et al., 2021) both have a left-handed stepping bias, as does the kinesin-5 Eg5 (Yajima et al., 2008), although this motor’s directional preference has recently been called into question (Meißner et al., 2023). By contrast, the minus-end-directed kinesin-14 Ncd (Mitra et al., 2020; Nitzsche et al., 2016; Walker et al., 1990) has a right-handed stepping bias, and the minus-end-directed cytoplasmic dynein also has a weak right-handed bias (Can et al., 2014; Elshenawy et al., 2019). These torques would be expected to additively twist the spindle in a left-handed direction. However, the human spindle exhibits only a weak left-handed twist on average (Neahring et al., 2021; Novak et al., 2018; Trupinic et al., 2022). It is not known how molecular-scale torques are balanced in the spindle to produce a relatively achiral structure from chiral motors.

The spindle’s left-handed twist was first quantified in metaphase HeLa and U2OS cells (Novak et al., 2018). Twist has been proposed to allow the metaphase spindle to accommodate mechanical load along the pole-to-pole axis (Trupinic et al., 2022), although its functional importance for chromosome segregation remains to be studied. Twist ranges from ∼0 to 2° of rotation per micron of displacement along the pole-pole axis depending on quantification method, cell type (with spindles in RPE1 cells having weaker twist than HeLa or U2OS spindles), and mitotic phase, peaking around anaphase onset (Trupinic et al., 2022). Several motors have been demonstrated to contribute to spindle twist in the predicted direction. Inhibiting Eg5, depleting the kinesin-8 KIF18A, or depleting the kinesin-6 MKLP1 reduces the spindle’s left-handed twist at metaphase in some cell types, suggesting that torques generated by biased motor stepping are relevant to the twist of the spindle as a whole (Novak et al., 2018; Trupinic et al., 2022). Only one perturbation has been demonstrated to increase the spindle’s left-handed twist: our previous work revealed that in anaphase RPE1 spindles, knockout of dynein’s targeting factor NuMA, combined with Eg5 inhibition to maintain spindle bipolarity, leads to strong left-handed twist (Neahring et al., 2021). Although it remains unknown how NuMA deletion increases spindle twist, the observation that twist can either be strengthened or abrogated by depleting various spindle factors raises the question of how opposing torques are generated and resisted to set spindle twist.

Here, we investigate how torques are balanced such that the spindle exhibits only slight global twist. We find that spindles in the human mammary epithelial cell line MCF10A exhibit stronger baseline twist than spindles in other cell lines studied to date, providing a system in which to study factors that both increase and decrease spindle twist. Using lattice light sheet microscopy, we show that twist is sustained at its strongest during anaphase, and we ask both how this twist is generated at anaphase and how it is restrained to prevent dramatic twist during chromosome segregation. The motors KIF4A and MKLP1, which redundantly contribute to spindle elongation at anaphase, are redundantly required for left-handed spindle twist, as is the actin cytoskeleton. Dynein and its cortical recruitment factor LGN counteract this twist. Together, our results show that force generators both within the spindle and at the cell periphery regulate twist in competing directions to set the spindle’s slight left-handed twist at anaphase.

## Results and discussion

### MCF10A spindles exhibit high baseline twist that peaks in late metaphase and anaphase

To study torque regulation in the spindle, we sought to identify a cell line in which spindles exhibited higher baseline twist than that observed in previously characterized cell lines. We reasoned that because twist differs between the human cell lines RPE1, HeLa, and U2OS (Neahring et al., 2021; Novak et al., 2018; Trupinic et al., 2022), other human cell lines may exhibit stronger twist, and that stronger twist would allow us greater dynamic range to study factors that both increase and decrease twist. We quantified twist using the optical flow method (Trupinic et al., 2022) in which we live-imaged full spindle volumes, computationally rotated the images to view the spindle along the pole-to-pole axis, and calculated the displacement fields of pixel intensities between successive frames from 30% and 70% of the pole-to-pole axis (Fig. 1 A). These flow vectors were converted to polar coordinates and averaged to produce a single twist value for each spindle. Compared to other methods of quantifying twist, such as manual bundle tracing (Neahring et al., 2021; Novak et al., 2018; Trupinic et al., 2022) or quantification based on bundle angles, the optical flow method is more strongly influenced by image noise but is more easily applied to large numbers of cells (Trupinic et al., 2022) (Fig. S1 A-B).

**Figure 1.**
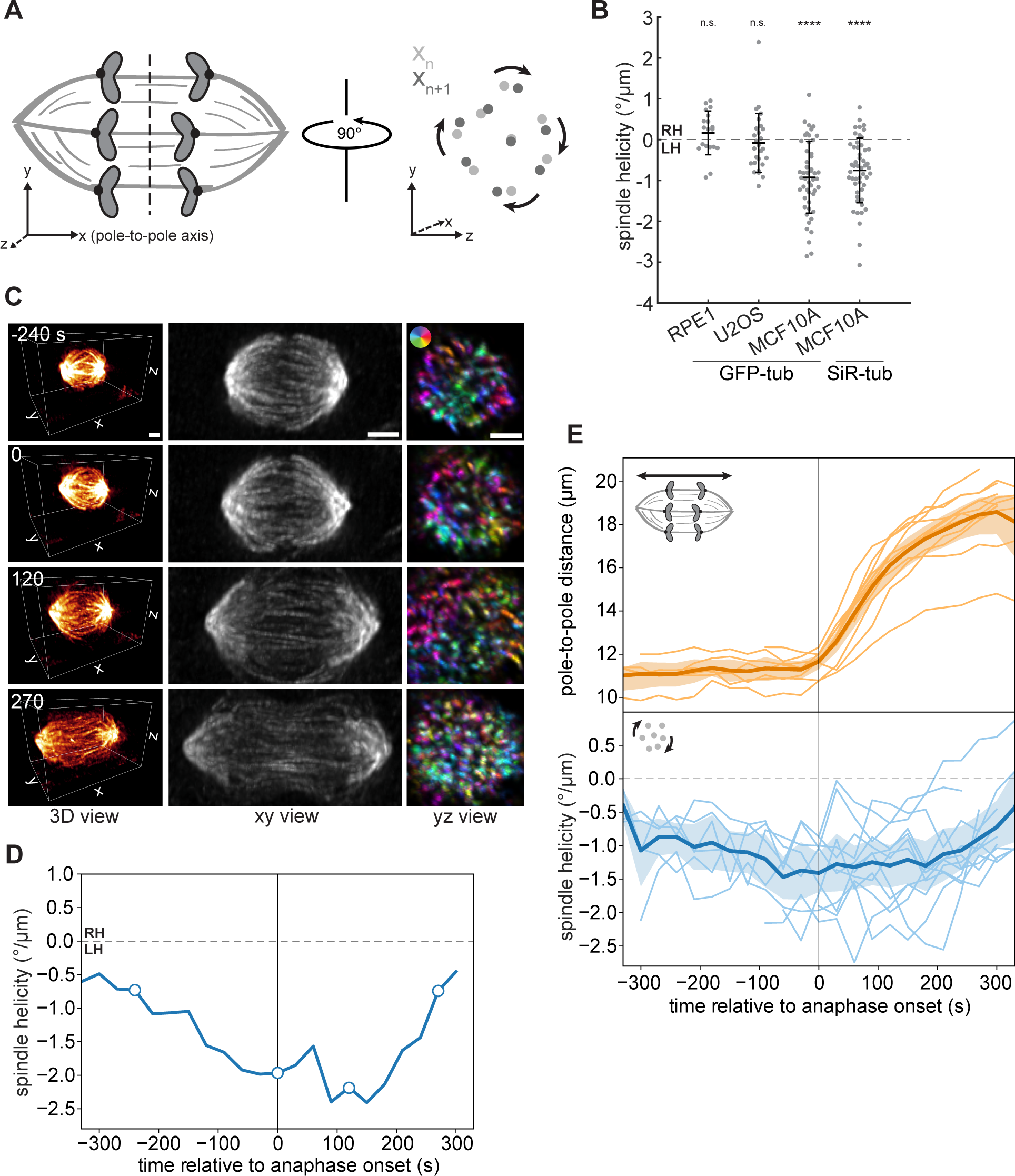
MCF10A spindles exhibit high baseline twist that peaks in late metaphase and anaphase. **(A)** Schematic diagram of spindle twist quantification, using the method developed by (Trupinic et al., 2022). Three-dimensional image stacks were rotated to view the spindle along the pole-to-pole axis. Farnebäck optical flow was computed between successive frames, and flow vectors were converted to polar coordinates and averaged for each spindle (see Materials and methods). **(B)** Spindle helicity (average degrees rotated around the pole-to-pole axis per µm displacement along the pole-to-pole axis) at anaphase in three human epithelial (RPE1, MCF10A) or epithelial-like (U2OS) cell lines, calculated from GFP-α-tubulin or SiR-tubulin intensity. Negative values represent left-handed helicity, and positive values represent right-handed helicity. Black lines represent mean ± SD. n = 19, 27, 50, and 51 spindles pooled from N = 2, 4, 5, and 5 independent experiments for RPE1 GFP-tub, U2OS GFP-tub, MCF10A GFP-tub, and MCF10A SiR-tub, respectively. n.s. not significant, ****p = 1.50×10^-9^ (MCF10A GFP-tub) and p = 1.11×10^-8^ (MCF10A SiR-tub), one-sample t-tests comparing each sample to a mean of 0. **(C)** Lattice light sheet images of the same MCF10A cell, labeled with SiR-tubulin, at four different timepoints. The xy view (center) shows maximum intensity projections of the entire spindle region. The yz view (right) shows maximum intensity projections between 30% and 70% of the pole-to-pole axis for the same image volumes after rotating them by 90°. Colors indicate directions of Farnebäck optical flow vectors, according to the color legend shown in the top image. Scale bars = 3 µm. **(D)** Helicity over time, calculated from lattice light sheet images, for the MCF10A spindle shown in (C). The four timepoints in (C) are indicated by open circles. **(E)** Length (upper panel) and helicity (lower panel) over time of 12 MCF10A spindles, calculated from time-lapse lattice light sheet images. The center line and shaded region represent the mean and 95% confidence interval.

Focusing on anaphase, when we previously observed that spindle twist is differentially regulated (Neahring et al., 2021), we found that the non-transformed mammary epithelial cell line MCF10A (Soule et al., 1990) exhibited strong spindle twist, with visually apparent left-handed twist in unperturbed cells. We quantified the twist of early- and mid-anaphase MCF10A cells labeled with either overexpressed GFP-tubulin or SiR-tubulin and found significant left-handed twist (negative helicity, mean -0.92 and -0.76°/µm respectively), while anaphase RPE1 and U2OS spindles were not significantly twisted (Fig. 1 B). Although metaphase U2OS spindles were previously reported to exhibit left-handed twist (Novak et al., 2018), our inability to detect twist here may be due to the difference in mitotic stage and/or to the reduced signal-to-noise ratio of the spindle midzone in spindles labeled with GFP-tubulin rather than GFP-PRC1.

Our comparison between different human cell lines was performed by live confocal imaging of a single timepoint per cell, leading us to wonder how twist changes during mitotic progression in MCF10A spindles. To image dividing cells volumetrically at high time resolution, we used lattice light sheet microscopy (Fig. 1 C-E). This imaging modality allowed us to obtain near-isotropic resolution with minimal phototoxicity, ideal for studying three-dimensional spindle architecture over time (Pamula et al., 2019). Comparing the profiles of 12 cells revealed several interesting features of spindle twist. Twist magnitude varied considerably from cell to cell, with peak helicities ranging from -1.26 to -2.74°/µm. The average twist across all cells in the first 4 minutes of anaphase was -1.28°/µm, consistent with the improved signal-to-noise ratio in these images compared to our confocal data. This highlights the fact that helicity magnitudes are not necessarily comparable between different imaging modalities or labeling methods, although twist is comparable within a given dataset (Fig. S1 A-B). On average, left-handed twist became stronger in the final minutes of metaphase, consistent with previous findings in HeLa and RPE1 cells (Trupinic et al., 2022). Twist was maintained throughout early and mid-anaphase for approximately 3 minutes, before dissipating in late anaphase, typically coinciding with a slower phase of spindle elongation (Fig. 1 C-E). Given the high baseline twist in MCF10A spindles, and given the sustained period of stronger twist in the first few minutes of anaphase, we focused thereafter on anaphase MCF10A cells to study how torques are generated and resisted in the spindle.

### The midzone motors KIF4A and MKLP1 are redundantly required for the anaphase spindle’s left-handed twist

We next asked what factors give rise to the spindle’s left-handed twist at anaphase. Although the motors Eg5 and KIF18A have been shown to promote left-handed twist at metaphase (Novak et al., 2018; Trupinic et al., 2022), many mitotic motors undergo changes in localization and function at anaphase, and the molecular basis of anaphase spindle twist has not been studied. Anaphase spindle elongation in human cells is powered by several kinesins that localize to antiparallel microtubule overlaps in midzone bundles, where they slide microtubules apart (Vukusic et al., 2017; Yu et al., 2019). KIF4A, Eg5, and the kinesin-6 motors MKLP1 and MKLP2 all redundantly contribute to spindle elongation (Fig. 2 A) (Vukusic et al., 2017; Vukusic et al., 2021), and we hypothesized that these kinesins may generate anaphase-specific left-handed torques.

**Figure 2.**
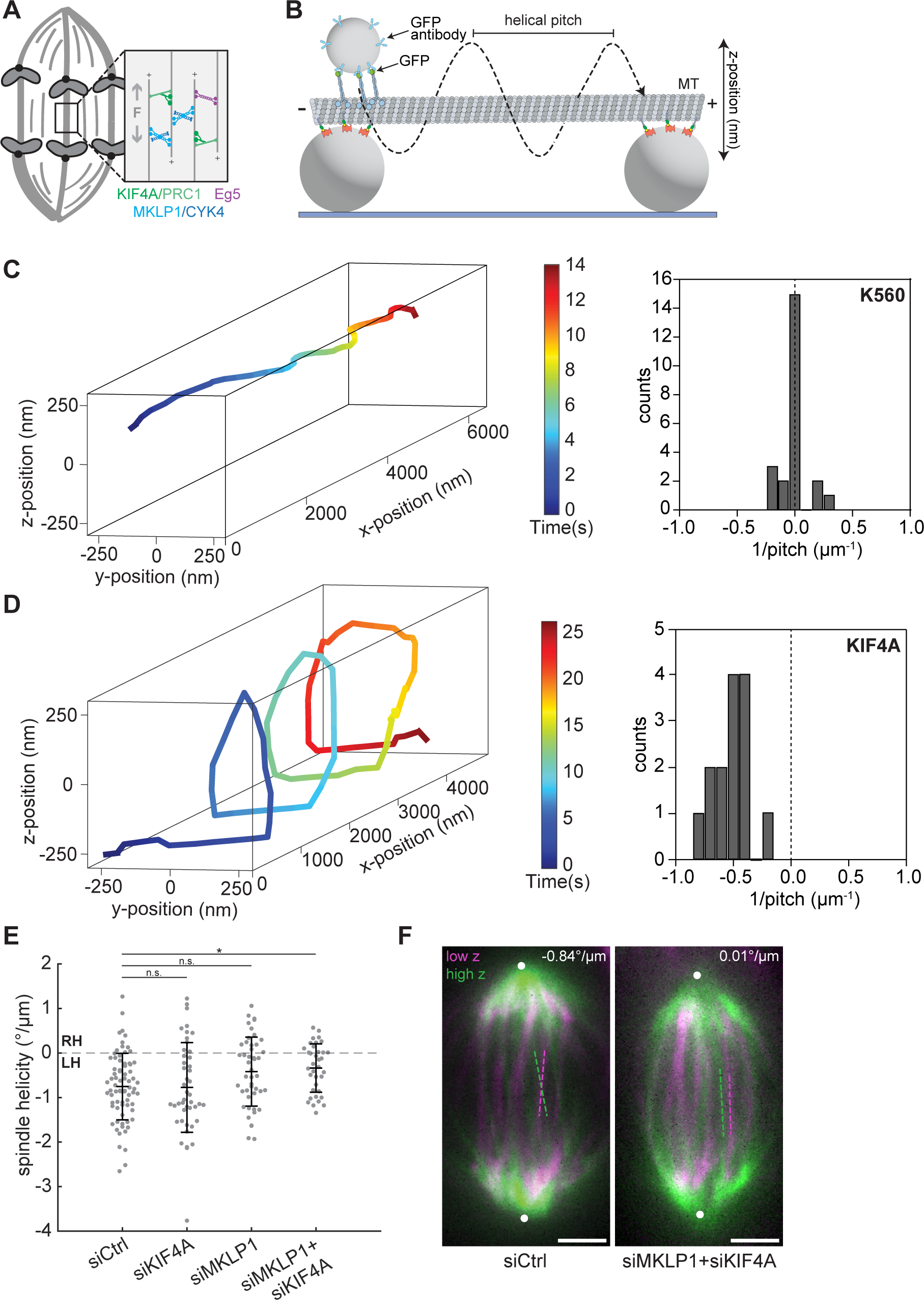
The midzone motors KIF4A and MKLP1 are redundantly required for the anaphase spindle’s left-handed twist. **(A)** Schematic diagram of the midzone motors KIF4A (green), MKLP1 (blue), and Eg5 (purple) that cooperate to drive anaphase spindle elongation (gray “F” and arrows). **(B)** Schematic diagram of the experimental geometry of the in vitro microtubule bridge assay (see Materials and methods, not to scale). A fluorescently labeled microtubule was suspended between two beads of 2 µm diameter. A 0.51 µm-diameter cargo bead was densely coated by multiple kinesins and brought onto the microtubule bridge by an optical trap (not shown) and bead motility was imaged using brightfield illumination. **(C)** (Left) Example 3D trajectory of a kinesin-1 (K560) coated cargo bead shows straight motility. (Right) Histogram of the inverse of the helical pitch of K560-coated beads (-0.004 ± 0.112 µm^-1^, mean ± SD, n = 23 rotations). The left-handed helical motion was defined as negative pitch. **(D)** (Left) Example 3D trajectory of a KIF4A-coated cargo bead shows left-handed helical motility. (Right) Histogram of the inverse of the helical pitch of KIF4A-coated beads (-0.53 ± 0.16 µm^-1^, n = 14 rotations). **(E)** Helicity of anaphase spindles calculated from SiR-tubulin intensity. Black lines represent mean ± SD. n = 74, 45, 46, and 36 spindles pooled from N = 7, 5, 7, and 6 independent experiments for siControl, siKIF4A, siMKLP1, and siMKLP+siKIF4A, respectively. n.s. not significant, *p = 0.048, one-way ANOVA with Tukey’s post-hoc test. **(F)** Confocal images of live MCF10A cells labeled with SiR-tubulin. Maximum intensity projections of a 2 µm-thick low region (magenta) and a 2 µm-thick high region (green) relative to the spindle midplane are overlaid. Dashed lines highlight individual microtubule bundles in each region. The helicity of each spindle is indicated in the top right. Positions of spindle poles (not visible in these high and low z-planes), manually assigned based on SiR-tubulin signal, are indicated by white circles. Scale bars = 3 µm.

Eg5 and the *C. elegans* homolog of MKLP1 have been shown to step in a left-handed fashion in vitro (Maruyama et al., 2021; Yajima et al., 2008), but the chirality of KIF4A has not been studied. Due to its ability to slide antiparallel microtubules apart (Hannabuss et al., 2019; Wijeratne and Subramanian, 2018), its anaphase-specific localization to midzone microtubule overlaps (Kurasawa et al., 2004), and its contribution to anaphase spindle elongation (Vukusic et al., 2021), KIF4A is a good candidate for a left-handed torque generator in the anaphase spindle. Thus, we characterized the torque generation of KIF4A on suspended microtubule bridges between 2 µm-diameter beads immobilized on a coverslip (Can et al., 2014) (Fig. 2 B). Smaller-sized (0.51 µm-diameter) cargo beads were decorated by multiple kinesin motors, brought close to the microtubule bridge with an optical trap, and their motility was tracked in three dimensions using brightfield microscopy (Fig. S2 A). We first used a truncated version of human kinesin-1 (KIF5B, amino acids 1-560, referred to as K560) to validate our experimental approach since K560 is known to follow a single protofilament on its microtubule track (Can et al., 2014; Ray et al., 1993; Yajima and Cross, 2005). We observed that cargo beads coated with K560 exhibited a combination of left-handed (6.6 ± 0.4 µm pitch; mean ± SD, 5 beads, 5 rotations), right-handed (4.7 ± 1.4 µm pitch; 3 beads, 3 rotations), and straight (15 beads) movements (Fig. 2 C), consistent with the reported pitch lengths of microtubules with 14, 12, or 13 protofilaments, respectively (Hyman et al., 1995; Ray et al., 1993). Unlike kinesin-1, all KIF4A-driven beads exhibited left-handed motility with a shorter pitch of 2.1 ± 0.7 µm (10 beads, 14 rotations; two-tailed t-test, p = 10^-4^; Fig. 2 D), demonstrating that KIF4A generates left-handed torque on microtubules.

We next sought to test whether the motors powering anaphase elongation contribute to left-handed spindle twist in their cellular context. We first tested the individual contributions of these motors by depleting KIF4A or MKLP1 or inhibiting Eg5 with S-trityl-L-cysteine (STLC). We confirmed that each cell analyzed displayed the expected late anaphase and cytokinesis phenotypes (for KIF4A and MKLP1 depletions; Fig. S2 B-F) or led to monopolar spindle formation in nearby prophase cells (for STLC treatment; data not shown). None of these perturbations significantly affected anaphase spindle twist (Fig. 2 E; Fig. S2 G). When we co-depleted KIF4A and MKLP1, however, spindles were significantly less twisted with a mean helicity of -0.34°/µm (Fig. 2 E and F; Fig. S2 D). This suggests that similarly to their redundant roles in elongating the anaphase spindle, the midzone motors KIF4A and MKLP1 redundantly generate left-handed torques to twist the anaphase spindle.

It is possible that spindle motors could modulate spindle twist by altering spindle shape or other microtubule properties, in addition to or instead of directly exerting torques on spindle microtubules. Indeed, a previous study noted that rounder HeLa spindles tended to exhibit stronger twist, although this correlation did not extend to the RPE1 cell line (Trupinic et al., 2022). We quantified anaphase spindle shape after depleting the motors KIF4A or MKLP1, and found that KIF4A knockdown was associated with slight changes in spindle shape: siKIF4A spindles were slightly wider, and had a lower length-to-width ratio, on average (Fig. S1 C-E). However, spindle helicity was not significantly correlated with spindle length, width, or aspect ratio (Fig. S1 F-H). These results suggest that KIF4A does not regulate spindle twist simply by changing the shape of the anaphase spindle, consistent with a functional role for its capacity to generate torque on microtubules.

### Actin contributes to the spindle’s left-handed twist at anaphase

Motor localization to the spindle midzone is only one of several changes that occurs at the metaphase-to-anaphase transition. Interactions between the spindle and the actin cytoskeleton also become more pronounced: NuMA-dynactin-dynein complexes enrich at the anaphase cell cortex in an actin-dependent manner (Kotak et al., 2013; Kotak et al., 2014), and the spindle midzone provides a spatial cue for actin assembly at the site of the future contractile ring (Pollard and O’Shaughnessy, 2019). Intriguingly, chiral flows of cortical actin break left-right symmetry by biasing anaphase spindle orientation in early embryos of *C. elegans* and the snail *Lymnaea stagnalis* (Davison et al., 2016; Kuroda et al., 2016; Naganathan et al., 2014; Shibazaki et al., 2004), and the actin cytoskeleton demonstrates intrinsic chirality in cultured human fibroblasts (Tee et al., 2015). Thus, we wondered whether chiral actin structures or flows could contribute to the human spindle’s chiral twist at anaphase.

We disrupted the actin cytoskeleton by treating MCF10A cells with latrunculin A (LatA), a small molecule that sequesters actin monomers, and found that this abrogated the anaphase spindle’s left-handed twist (mean -0.19°/µm, compared to -0.71°/µm after DMSO treatment). By contrast, treating cells with the myosin II inhibitor blebbistatin or the ROCK inhibitor Y27632 did not affect spindle twist (Fig. 3 A and B). We confirmed that these drugs disrupted actomyosin contractility by imaging cells later in anaphase, when they blocked cytokinetic furrow ingression (Fig. 3 C). These results indicate that the actin cytoskeleton, but not actomyosin contractility, reinforces the anaphase spindle’s left-handed twist. Thus, the contribution of actin to human spindle twist is distinct from the myosin-dependent cortical flows that influence cellular chirality in some invertebrate embryos.

**Figure 3.**
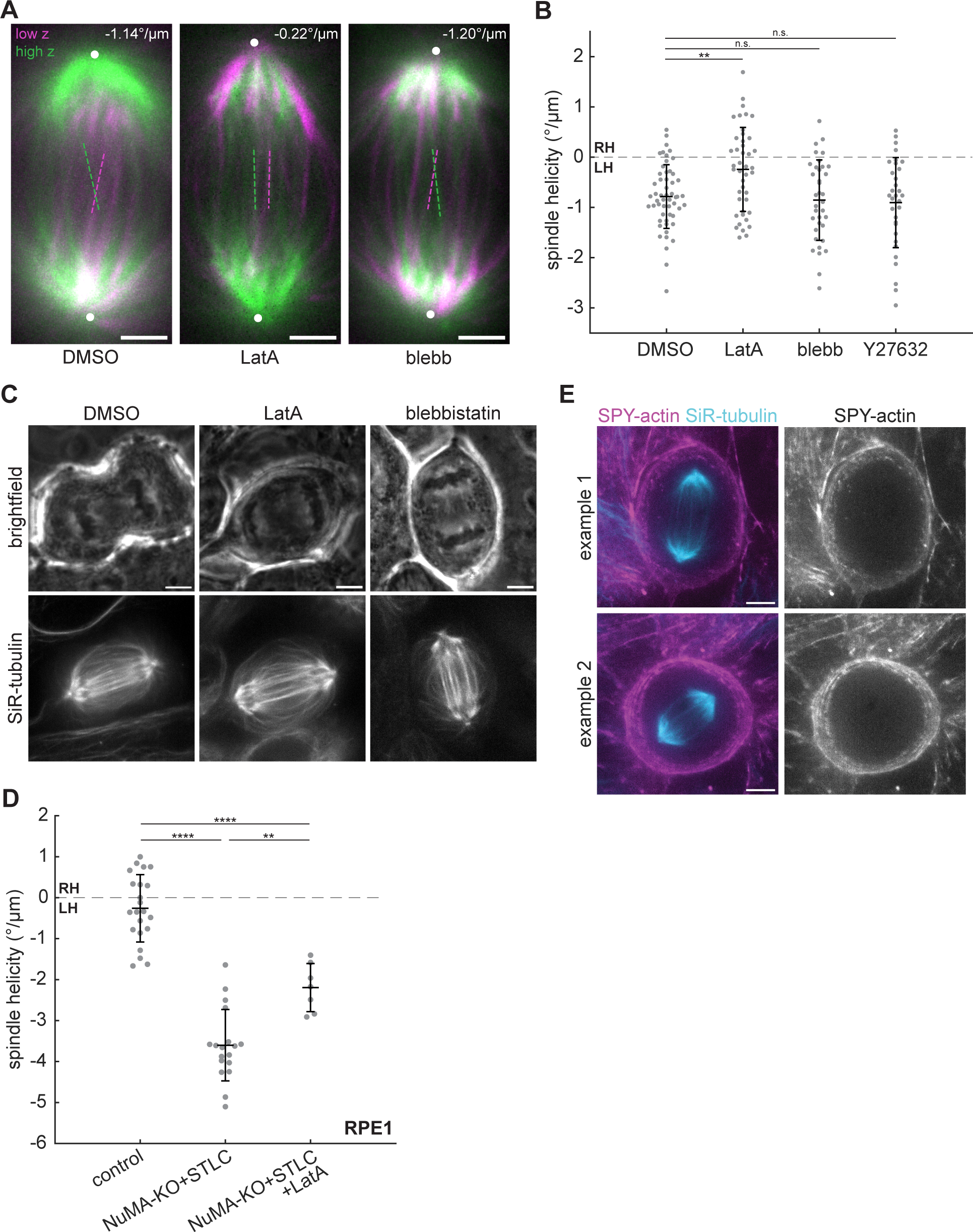
Actin contributes to the spindle’s left-handed twist at anaphase. **(A)** Confocal images of live MCF10A cells treated with 0.1% DMSO, 500 nM latrunculin A, or 25 µM blebbistatin labeled with SiR-tubulin. Maximum intensity projections of a 2 µm-thick low region (magenta) and a 2 µm-thick high region (green) relative to the spindle midplane are overlaid. Dashed lines highlight individual microtubule bundles in each region. The helicity of each spindle is indicated in the top right. Positions of spindle poles are indicated by white circles. Scale bars = 3 µm. **(B)** Helicity of anaphase spindles calculated from SiR-tubulin intensity. Black lines represent mean ± SD. n = 54, 43, 36, and 32 spindles pooled from N = 6, 5, 4, and 3 independent experiments for DMSO, LatA, blebbistatin, and Y27632, respectively. n.s. not significant, **p = 0.0039, one-way ANOVA with Tukey’s post-hoc test. **(C)** Confocal images of late anaphase MCF10A cells. 500 nM LatA and 25 µM blebbistatin treatment each block cytokinetic furrow ingression. Brightfield images (upper row) represent a single z-plane and SiR-tubulin images (lower row) are maximum intensity projections of 10 µm z-stacks. **(D)** Helicity of anaphase RPE1 spindles, synchronized with RO-3306, calculated from GFP-tubulin intensity. The control and NuMA-KO+STLC conditions are the same cells included in a previous publication (Neahring et al., 2021), re-analyzed using the optical flow method. Black lines represent mean ± SD. n = 22, 18, and 7 spindles pooled from N = 4, 5, and 3 independent experiments for control, NuMA-KO+STLC, and NuMA-KO+STLC+LatA, respectively. **p = 9.99×10^-4^, ****p = 4.32×10^-16^ (control vs. NuMA-KO+STLC) and ****p = 5.93×10^-6^ (control vs. NuMA-KO+STLC+LatA). **(E)** Live confocal images (maximum intensity projections of 10 µm z-stacks) of control anaphase MCF10A cells labeled with SPY555-actin (magenta) and SiR-tubulin (cyan). The SPY555-actin channel is shown alone at right. Scale bars = 5 µm.

To test the generality of actin’s contribution to spindle twist, we tested its role in a different experimental model. We previously found that normally achiral RPE1 spindles are strongly left-handed in anaphase when NuMA and Eg5 are co-inhibited (Neahring et al., 2021). Treating these doubly inhibited RPE1 cells with LatA also significantly reduced spindle twist by 39% (Fig. 3 D). Although LatA has been reported to have no effect on the twist of metaphase HeLa spindles (Novak et al., 2018), the discrepancy with our results in anaphase MCF10A and RPE1 cells could be due to actin’s anaphase-specific roles.

To gain further insight into the actin cytoskeleton’s influence on spindle twist, we live-imaged the localization of actin in dividing MCF10A cells labeled with SPY555-actin. Although actin filaments accumulate around early anaphase centrosomes in some cell types (Farina et al., 2019), we could not detect centrosome-localized actin in MCF10A cells, and instead observed SPY555-actin signal only at the cell periphery (Fig. 3 E). Surprisingly, these observations suggest that factors not just within the spindle body, but also actin at the cell periphery, regulate spindle twist.

### Dynein counteracts left-handed twist in the anaphase spindle

The motor-generated torques characterized to date are expected to additively twist the spindle, and the effects of the actin cytoskeleton described above contribute to twist in the same left-handed direction (Fig. 3). Thus, we sought to understand what factors are required to oppose left-handed torques so that the spindle exhibits only slight global twist. Our previous work identified NuMA and Eg5 co-inhibition in RPE1 spindles as the first known perturbation that increases the spindle’s left-handed twist (Neahring et al., 2021). Since inhibiting Eg5 alone had no effect in that system, we hypothesized that NuMA and its interactors dynactin and dynein (Fig. 4 A) play key roles in resisting spindle twist. Consistent with this hypothesis, siRNA-mediated depletion of dynein heavy chain indeed increased left-handed twist in anaphase MCF10A spindles (Fig. 4 B and C; Fig. S3 A).

**Figure 4.**
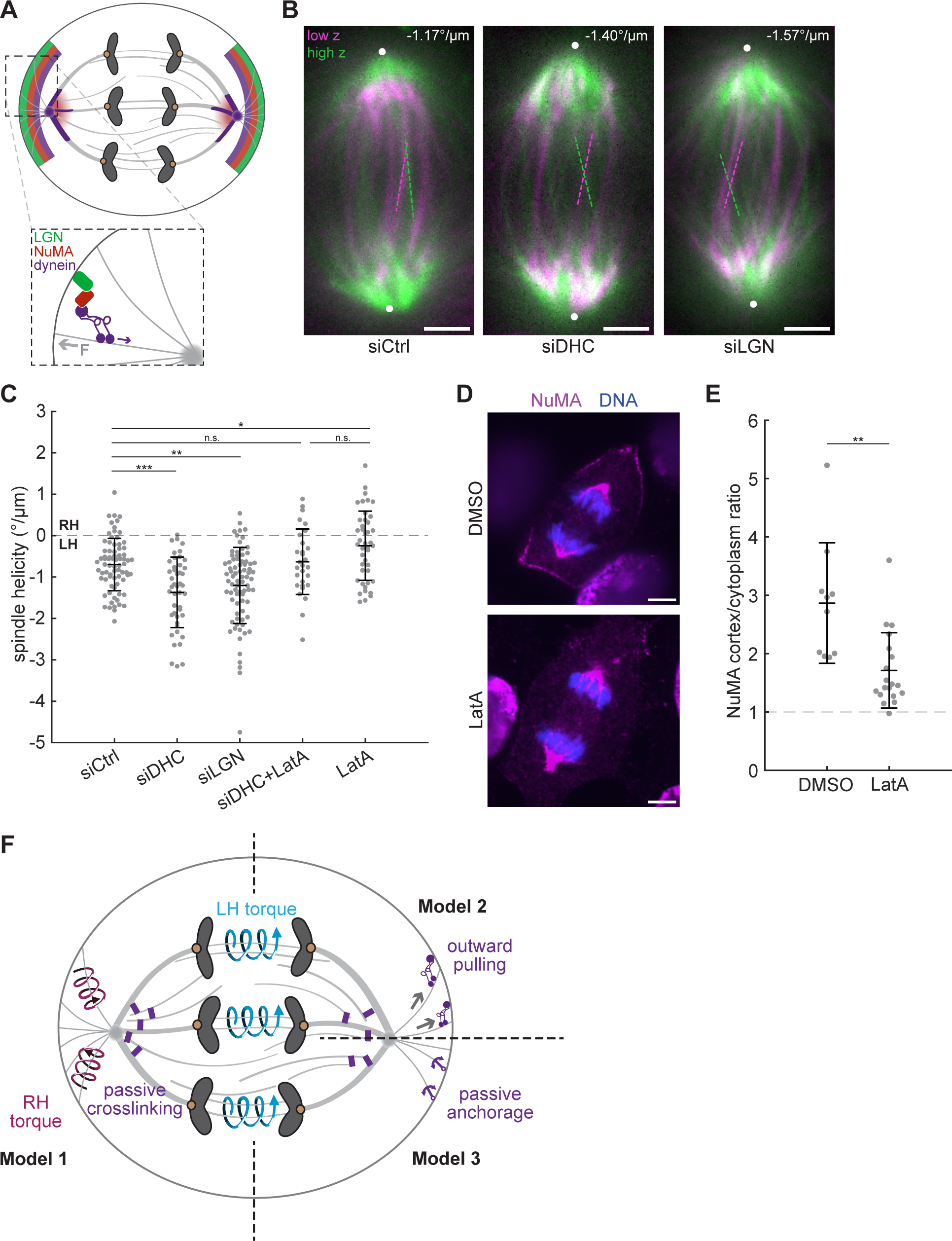
Dynein counteracts left-handed twist in the anaphase spindle. **(A)** Schematic diagram of LGN, NuMA, and dynein localization in anaphase cells. Dynein and NuMA cluster microtubule minus ends and localize to spindle poles, while LGN-NuMA-dynein complexes localize to cortical crescents where they exert pulling forces on astral microtubules. The purple arrow indicates direction of dynein stepping, and the gray arrow indicates direction of force on astral microtubules. **(B)** Confocal images of live MCF10A cells transfected with siRNA targeting luciferase (siCtrl), dynein heavy chain, or LGN labeled with SiR-tubulin. Maximum intensity projections of a 2 µm-thick low region (magenta) and a 2 µm-thick high region (green) relative to the spindle midplane are overlaid. Dashed lines highlight individual microtubule bundles in each region. The helicity of each spindle is indicated in the top right. Positions of spindle poles are indicated by white circles. Scale bars = 3 µm. **(C)** Helicity of anaphase spindles calculated from SiR-tubulin intensity. Black lines represent mean ± SD. n = 75, 45, 78, 29, and 43 spindles pooled from N = 7, 5, 7, 3, and 5 independent experiments for siCtrl, siDHC, siLGN, siDHC+LatA, and LatA, respectively. n.s. not significant, *p = 0.027, **p = 0.0011, ***p = 1.062×10^-4^, one-way ANOVA with Tukey’s post-hoc test. LatA data represents the same cells shown in Fig. 3. **(D)** Immunofluorescence images (single z-planes) of anaphase MCF10A cells stained for NuMA (magenta) and DNA (blue). Scale bars = 5 µm. **(E)** Quantification of NuMA enrichment at the cell cortex relative to NuMA intensity in the cytoplasm (see Materials and methods), in immunofluorescence images of MCF10A cells treated with DMSO or LatA. Dashed line indicates a cortex/cytoplasm ratio of 1, i.e. no cortical enrichment. Black lines represent mean ± SD. n = 10 cells from 1 experiment (DMSO), 19 cells from 2 experiments (LatA). **p = 9.47×10^-4^, two-sample t-test. **(F)** Proposed models for twist regulation in the anaphase spindle. Motors within the spindle midzone generate left-handed torques (blue arrows), which are partially counteracted by passive dynein-mediated crosslinking at spindle poles (purple rectangles) and by LGN-NuMA-dynein complexes at the cell cortex. These cortical complexes could actively generate right-handed torques on astral microtubules (magenta arrows, Model 1), exert outward pulling on the spindle poles (Model 2), or serve to anchor astral microtubules in the cortex (purple anchors, Model 3). Together, these active and passive torques establish weak global left-handed twist in the anaphase spindle.

We next probed the mechanism by which dynein counteracts left-handed twist. At mitosis NuMA recruits dynactin and dynein to microtubule minus ends, where they act as a complex to cluster minus ends at spindle poles (Gaglio et al., 1996; Heald et al., 1996; Hueschen et al., 2017; Merdes et al., 1996; Verde et al., 1991). NuMA-dynactin-dynein complexes also localize to the cell cortex, where they generate pulling forces on astral microtubules to position the spindle (Kotak et al., 2012) (Fig. 4 A). Based on mammalian dynein’s in vitro stepping behavior with other adaptors (Elshenawy et al., 2019), its torque generation inside the spindle would be predicted to augment the spindle’s left-handed twist. Because we observed the opposite phenotype upon dynein depletion, we reasoned that dynein likely does not regulate spindle twist by generating torques between spindle microtubule pairs. Instead, we considered two alternative explanations: first, the cortical pool of NuMA-dynactin-dynein could resist left-handed spindle twist, or alternatively that the stiff, microtubule-dense spindle poles built by NuMA-dynactin-dynein activity could non-directionally resist torques generated by other motors.

To test the cortical pool of dynein, we depleted LGN, one of several factors that recruits NuMA-dynactin-dynein complexes to the cortex during anaphase (Fig. S3 B) (Du and Macara, 2004; Kiyomitsu and Cheeseman, 2013; Kotak et al., 2014; Seldin et al., 2013). We confirmed that upon LGN depletion, NuMA’s cortical localization was reduced in anaphase MCF10A cells (Fig. S3 C and D). LGN knockdown increased the spindle’s left-handed twist to an average of -1.20°/µm, significantly stronger than that of control-depleted cells and almost as strong as that of dynein-depleted cells (Fig. 4 B and C). This suggests that the cortical pool of NuMA-dynactin-dynein indeed counteracts left-handed twist in the anaphase spindle.

Finally, we tested whether the pool of dynein within the spindle plays a role in resisting spindle twist. To do so, we combined dynein depletion with LatA treatment, a perturbation that reduces NuMA-dynactin-dynein localization to the cortex (Fig. 4 D and E) (Kotak et al., 2014). By comparing twist in LatA-treated versus dynein-depleted and LatA-treated cells—both conditions in which little NuMA/dynein is present at the cortex—we could probe the contribution of spindle-localized dynein. Compared to LatA treatment alone, dynein depletion combined with LatA treatment resulted in increased left-handed twist, although this result did not reach statistical significance (Fig. 4 C, mean = -0.24 and -0.59°/µm). Similarly to our analysis of spindle shape after depleting the anaphase motors KIF4A and MKLP1, we quantified spindle shape after LatA treatment and depletion of dynein or LGN. While each of these treatments affected overall spindle shape, helicity was uncorrelated with spindle length, width, and aspect ratio (Fig. S1 C-H), suggesting that altered spindle dimensions do not mediate the effects of these perturbations. Taken together, we conclude that the cortical pool of dynein counteracts left-handed twist, and our data suggests that the spindle pool of dynein also contributes to this effect. Thus, dynein is required to prevent strong global left-handed twist.

In this study, we investigate how the spindle attains its relatively untwisted architecture despite being built by chiral force-generators. Focusing on anaphase, when we find that the spindle’s strongest twist is sustained for several minutes (Fig. 1), we identify factors that promote left-handed twist (Figs. 2 and 3) and that counteract it (Fig. 4). Unexpectedly, we find that twist is regulated not just by motors internal to the spindle, but also by microtubule-associated proteins and actin at the cell periphery. Together, our results demonstrate that spindle twist is an emergent phenomenon that integrates inputs from the spindle’s broader cellular environment.

Motors that crosslink, slide, and twist microtubules are abundant in the spindle, and have been the focus of most work on spindle twist to date. We find that the midzone motors KIF4A and MKLP1 are redundantly required for left-handed twist, similar to their redundant contributions to spindle elongation (Vukusic et al., 2021), reflecting a common design principle in the anaphase spindle. Both KIF4A and MKLP1 can slide antiparallel microtubules apart in vitro (Hannabuss et al., 2019; Nislow et al., 1992; Wijeratne and Subramanian, 2018), both step in a left-handed direction around microtubules (Fig. 2) (Maruyama et al., 2021), and both localize and concentrate at the midzone at anaphase onset (Kurasawa et al., 2004; Matuliene and Kuriyama, 2002), suggesting that they may increase twist at anaphase by directly exerting torques on overlapping antiparallel microtubules. However, we cannot exclude that KIF4A and/or MKLP1 could regulate twist indirectly via effects on the midzone’s microtubule dynamics or organization, potentially altering its material properties or the localization of downstream factors. Directly linking motor-generated torques to global spindle twist awaits the development of mutant motors with altered torque-generating capacities, an exciting future direction.

Opposing the effects of KIF4A and MKLP1, dynein and its targeting factors NuMA and LGN are each required to restrain the anaphase spindle’s left-handed twist. Although another study found that dynein inhibition did not increase twist in metaphase HeLa or RPE1 spindles (Trupinic et al., 2022), we observe strong phenotypes after dynein depletion in MCF10A cells or NuMA knockout in RPE1 cells (Fig. 4). Our results may differ due to our NuMA/dynein inhibition strategies or because we focus on the anaphase spindle, when NuMA and dynein partition more strongly to the cortex. We find evidence that the cortical pool of dynein counteracts the spindle’s left-handed twist, and that the spindle pool exhibits a trend in the same direction. LGN-NuMA-dynein complexes at the cortex could reduce twist by exerting active right-handed torques on astral microtubules (Fig. 4 F, Model 1), by pulling outwards on astral microtubules and spindle poles (Fig. 4F, Model 2), or by increasing astral microtubules’ anchorage in the cortex (Fig. 4 F, Model 3). NuMA and dynein are strongly enriched at spindle poles, and we propose that this pool could resist spindle twist by crosslinking microtubule minus ends to neighboring microtubules and by organizing stiff spindle poles. Although dynein has a slight right-handed stepping bias in vitro (Can et al., 2014; Elshenawy et al., 2019), this directional preference is inconsistent with the spindle-scale phenotypes we observe if dynein were to act primarily by generating torques between microtubule pairs. Dynein and its cofactors illustrate that motors can regulate spindle twist in many ways— by crosslinking microtubules, shaping spindle poles, and mediating interactions between the spindle and cell periphery—beyond twisting spindle microtubules around each other.

Finally, we find that the actin cytoskeleton is required for the anaphase spindle’s left-handed twist, both in MCF10A cells and in NuMA- and Eg5-inhibited RPE1 cells (Fig. 3). We cannot attribute the effects of actin on spindle twist to cortical flows or cortical stiffness, because inhibiting actomyosin contractility had no effect on twist. The role of actin is also independent from that of the LGN-NuMA-dynein cortical force-generating machinery, since they influence spindle twist in opposite directions. We speculate that the actin cytoskeleton may regulate spindle twist by controlling overall cell shape. In the future, further molecular dissection of actin-related proteins may provide insight into the role of actin uncovered here, since many myosins, formins, and actin filaments themselves are intrinsically chiral (Ali et al., 2002; Depue and Rice, 1965; Lebreton et al., 2018; Mizuno et al., 2011). Although the underlying mechanisms are not yet clear, our finding that the actin cytoskeleton affects twist is exciting because it reveals that multiple cytoskeletal systems coordinately regulate spindle twist.

In conclusion, our study shows that the human anaphase spindle’s weak left-handed twist requires both left-handed torque generators and factors that oppose them. The study of spindle twist is a recent area of inquiry, and many open questions remain. For example, it is unclear why twist differs between different cell types and between species (Velle et al., 2022), or whether twist is affected by other classes of mechanism such as microtubule crosslinking, the spindle’s material properties (Forth and Kapoor, 2017) including its torsional rigidity, or the turnover rates of microtubules and microtubule-associated proteins (Asthana et al., 2021; do Rosario et al., 2023). Finally, it will be interesting to explore potential functions of spindle twist in future work: are there adverse consequences for chromosome segregation if the spindle is too twisted, or not twisted enough? Twist could provide a mechanical advantage to the anaphase spindle as it separates sister chromosomes and initiates elongation, but conversely, our previous work has shown that strongly twisted NuMA- and Eg5-inhibited RPE1 spindles have high rates of chromosome segregation errors (Neahring et al., 2021). More broadly, our work motivates the study of how other cellular structures built from chiral molecular components either co-opt this chirality for their physiological function (for example, chiral actin flows in left-right symmetry breaking) (Naganathan et al., 2016), or balance chiral elements to restrain asymmetry.

## Materials and methods

### Cell culture

U2OS cells (female human osteosarcoma cells) were a gift from Samara Reck-Peterson (University of California San Diego, San Diego, CA, USA), and hTERT-RPE1 cells (female human retinal epithelial cells) were a gift from Bo Huang (University of California San Francisco, San Francisco, CA, USA). Both cell lines were cultured in DMEM/F12 (Thermo Fisher 11320) supplemented with 10% fetal bovine serum (Gibco 10438026). MCF10A cells (female human mammary epithelial cells) were purchased from ATCC (CRL-10317) and cultured as recommended by ATCC in MEGM (Lonza CC-3150) supplemented with bovine pituitary extract, insulin, hydrocortisone, and human epidermal growth factor according to the manufacturer’s instructions, and 100 ng/ml cholera toxin (Sigma-Aldrich C8052). Inducible NuMA-KO RPE1 cells stably expressing GFP-tubulin and mCherry-H2B (Neahring et al., 2021) were grown in DMEM/F12 supplemented with 10% tetracycline-screened FBS (Peak Serum PS-FB2). SpCas9 expression was induced by adding 1 µg/ml doxycycline hyclate 4 days before each experiment, refreshed after 24h and 48h. All cells were maintained at 37° and 5% CO_2_.

### Transfection, dyes, and small molecule treatments

For siRNA knockdowns, cells were transfected with siRNA targeting luciferase as a negative control (5’-CGUACGCGGAAUACUUCGA-3’, 50 pmol), LGN (Dharmacon ON-TARGETplus SMARTpool, L-004092-00-0005, 100 pmol), dynein heavy chain (5’-AAGGATCAAACATGACGGAAT-3’, 50 pmol) (Draviam et al., 2006; Tanenbaum et al., 2008), MKLP1 (pool of 3 sequences, Santa Cruz Biotechnology sc-35936, 50 pmol), or siKIF4A (pool of 3 sequences, Santa Cruz Biotechnology sc-60888, 50 pmol) for 48 hours using Lipofectamine RNAiMAX (Thermo Fisher 13778075) according to the manufacturer’s recommendations.

GFP-α-tubulin was expressed in RPE1, U2OS, and MCF10A cells by infection with BacMam virus. The GFP-α-tubulin coding sequence was cloned into the pEG BacMam vector (a gift from Eric Gouaux, Addgene plasmid #160451), recombinant bacmid DNA was generated in DH10Bac cells (Thermo Fisher 10361012), and isolated bacmid DNA was transfected into Sf9 cells (a gift from Yifan Cheng, University of California San Francisco, San Francisco, CA, USA) using Cellfectin II (Thermo Fisher 10362100) for production and amplification of BacMam virus according to a previously described protocol (Goehring et al., 2014). P2 BacMam virus was added to cells 2 days prior to imaging. Alternatively, tubulin was labeled by adding 100 nM SiR-tubulin and 10 µM verapamil for 30-60 minutes prior to imaging (Cytoskeleton, Inc. CY-SC002). Actin was labeled by diluting SPY555-actin (Cytoskeleton, Inc. CY-SC202) 1:1000 in media and incubating for 60 minutes prior to imaging.

For acute drug treatments, latrunculin A was added to a final concentration of 500 nM for 20 minutes prior to imaging, blebbistatin was added to a final concentration of 25 µM for 30 minutes prior to imaging, Y27632 was added to a final concentration of 10 µM for 30 minutes prior to imaging, STLC was added to a final concentration of 10 µM for 15 minutes prior to imaging, cytochalasin D was added to a final concentration of 5 µg/ml for 30 minutes prior to imaging, and DMSO was added to a final concentration of 0.1% (v/v) for 30 minutes prior to imaging.

For experiments in the RPE1 inducible NuMA-KO cell line (Fig. 3 D), cells were synchronized at the G2/M checkpoint by incubation overnight in 9 µM of the CDK1 inhibitor RO-3306. Cells were released into mitosis by washing 4X in warm media, and were imaged from prometaphase (approximately 30 minutes in controls) or after reaching the turbulent state (to confirm NuMA knockout; approximately 60 minutes in +doxycycline NuMA-KO cells). STLC (Sigma) and/or latruculin A (Invitrogen) were added to final concentrations of 5 µM and 500 nM respectively. Cells that entered anaphase within 90 minutes of drug addition were used for analysis.

### Western blotting

Cells grown in 6-well plates were lysed, and protein extracts were collected after centrifugation at 4°C for 30 min. Protein concentrations were measured using a Bradford assay kit (Bio-Rad), and equal concentrations of each sample were separated on 4-12% Bis-Tris gels (Invitrogen) by SDS-PAGE and transferred to a nitrocellulose membrane. Membranes were blocked with 4% milk in TBST (tris-buffered saline + 0.1% Tween 20), incubated in primary antibodies overnight at 4°C, and incubated with HRP-conjugated secondary antibodies for 45 minutes. Proteins were detected using SuperSignal West Pico or Femto chemiluminescent substrates. The following primary antibodies were used: mouse monoclonal anti-GAPDH (1:1,000, clone 258, Thermo Fisher 437000, RRID:AB_2532218), rabbit anti-KIF4A (1:1,000, Bethyl A301-074A, RRID:AB_2280904), mouse anti-MKLP1 C-12 (1:50, Santa Cruz Biotechnology sc-390113, RRID:AB_2802172), rabbit anti-LGN (1:1,000, Bethyl A303-032A, RRID:AB_10749181), and mouse monoclonal anti-dynein intermediate chain (1:500, clone 74.1, MilliporeSigma MAB1618, RRID:AB_2246059). The following secondary antibodies were used at a 1:10,000 dilution: mouse anti-rabbit IgG-HRP (Santa Cruz Biotechnology sc-2357, RRID:AB_628497) and mouse IgGκ BP-HRP (Santa Cruz Biotechnology sc-516102, RRID:AB_2687626).

### Immunofluorescence

Cells were plated on acid-cleaned, poly-L-lysine-coated, #1.5 25 mm coverslips for 3 days. Coverslips were washed in phosphate-buffered saline (PBS), fixed in MeOH pre-chilled to -20°C for 3 minutes, and washed again in PBS. Coverslips were blocked in TBST (0.05% Triton-X-100 in tris-buffered saline) containing 2% (w/v) bovine serum albumin. Antibodies were diluted in TBST containing 2% BSA and incubated for 1 hour (primary antibodies) or 45 minutes (secondary antibodies) at room temperature, followed by 4 washes in TBST. DNA was labeled with 1 µg/ml Hoechst 33342 prior to mounting on slides with ProLong Gold Antifade Mountant (Thermo Fisher). The following primary antibodies were used: rabbit anti-NuMA (1:300, NB500-174, Novus Biologicals; RRID:AB_10002562) and rat anti-α-tubulin (1:2,000, MCA77G, Bio-Rad; RRID:AB_325003). The following secondary antibodies were used at a 1:400 dilution: goat anti-rabbit AlexaFluor 568 and AlexaFluor 647 (A-11011 and A-21244, Thermo Fisher; RRID:AB_143157 and RRID:AB_2535812), goat anti-rat AlexaFluor 488 and AlexaFluor 647 (A-11006 and A-21247, Thermo Fisher; RRID:AB_2534074 and RRID:AB_141778). Brightness/contrast for each channel were scaled identically within each immunofluorescence experiment shown.

### Confocal microscopy

Cells were plated onto #1.5 glass-bottom 35 mm dishes coated with poly-D-lysine (MatTek Life Sciences P35G-1.5-20-C) 2-3 days prior to imaging and imaged in a humidified stage-top incubator maintained at 37° and 5% CO_2_ (Tokai Hit). Cells were imaged on a spinning disk (CSU-X1, Yokogawa) confocal inverted microscope (Eclipse Ti-E, Nikon Instruments) with the following components: 100× 1.45 NA Ph3 oil objective (Nikon); Di01-T405/488/568/647 head dichroic (Semrock); 405 nm (100 mW), 488 nm (150 mW), 561 nm (100 mW) and 642 nm (100 mW) diode lasers; ET455/50M, ET525/50M, ET630/75M, and ET690/50M emission filters (Chroma Technology); and a Zyla 4.2 sCMOS camera (Andor Technology).

### Lattice light sheet microscopy

The LLSM was a modified version of the microscope described in (Liu et al., 2018) and was controlled with custom LabVIEW software licensed from Janelia Research Campus, HHMI. Cells were plated on round 25 mm coverslips (Thorlabs CG15XH) coated with 200 nm fluorescent beads (Invitrogen FluoSpheres Carboxylate-Modified Microspheres, Ex/Em 660/680, F8807) to measure point spread functions for deconvolution and to align the lattice light sheet. The coverslip, excitation objective (Thorlabs water dipping objective, 0.60 NA, TL20X-MPL), and detection objective (Zeiss water dipping objective, 1.0 NA, 421452-9800-000) were immersed in approximately 50 ml of phenol red-free MCF10A culture medium maintained at 37°C and 5% CO_2_. Cells were labeled with 100 nM SiR-tubulin and 10 µM verapamil, and imaged using a 642 nm laser (MPB Communications Inc 2RU-VFL-P-2000-642-B1R) operating with 200 µW input power at the back pupil of the excitation objective. The 3D volumes consisting of 100 planes were acquired with 50 ms exposure per plane and iterating every 30 seconds by scanning the sample stage (SmarAct MLS-3252 Electromagnetic Direct-Drive) with a 400 nm step (corresponding to ∼215 nm along the optical z axis). A dithered harmonic-balanced hexagonal lattice light sheet (LLS) pattern with a numerical aperture of 0.35 and a sigma value of 0.09 was used (Liu et al., 2023). This LLS yields a 3D theoretical resolution of 340 x 340 x 570 nm at the 680 nm emission, which ensures that visualization and analysis of 3D spindle dynamics are not compromised by the axis of poorest resolution. A mask with an aperture of NA_min_ 0.3/NA_max_ 0.4 was used to block the unmodulated light and the higher order diffractions. Emission light was filtered by a Semrock FF02-685/40-25 band pass filter and captured by a Hamamatsu ORCA-Fusion sCMOS camera. Images were deconvolved, deskewed and rotated on a high-performance computing cluster using code available at https://github.com/abcucberkeley/LLSM5DTools/.

### Suspended microtubule bridge assay

#### Protein preparation

The truncated human kinesin-1 (KIF5B, amino acids 1-560) was expressed and purified using the baculovirus expression system in insect cells. GFP was fused to the C-terminus of kinesin to link motors to the antibody-coated beads. The recombinant full-length human KIF4A-GFP was expressed and purified in Sf9 cells as described (Subramanian et al., 2013). Microtubules were polymerized in BRB80 buffer using a mixture of LD655-labeled tubulin and unlabeled pig brain tubulin at a 1:4 ratio in the presence of 100 μM taxol.

#### Labeling cargo beads with antibodies

Carboxyl latex beads (Invitrogen) with a diameter of 0.51 μm were coated with anti-GFP antibodies as previously described (Belyy et al., 2014). The beads were initially pelleted and resuspended in activation buffer (10mM MES, 100 mM NaCl, pH 6.0). The carboxyl groups on the bead surfaces were functionalized with amine-reactive groups via *N-* ethyl-*N’-*(3-(dimethylamino)propyl)carbodiimide (EDC) and sulfo-*N-*hydroxysuccinimide (sulfo-NHS, Thermo Fisher) crosslinking for 20 minutes at 4°C. Extra crosslinking reagents were then removed by centrifugation, and beads were resuspended in PBS at pH 7.4. Anti-GFP polyclonal antibodies (Covance CA 5314/15, custom produced) were added to the beads and incubated at room temperature for 20 minutes. The surface of the beads was passivated by incubation with 5 mg/ml bovine serum albumin (BSA) overnight at 4°C. Excess antibodies were removed by centrifugation. Beads were then resuspended in PBS with 0.1% azide for storage at 4°C.

#### Preparation of flow chambers

Custom PEG-biotin coverslips were prepared as previously described (Zhao et al., 2023). Streptavidin-coated beads (2 μm diameter, Spherotech) were introduced into the chamber and incubated for 3 minutes to allow the beads to settle on the biotinylated coverslip surface. The chamber was washed with 15 µL motility buffer (MB; 30 mM HEPES, 5 mM MgSO_4_, 1 mM EGTA, pH 7.4 with KOH) and then incubated with 20 µL of 0.65 µM biotinylated chimeric protein in which seryl tRNA synthetase (SRS) was fused to a dynein microtubule binding domain (Dynein-SRS_85:82_ MTBD), which stably binds to microtubules without generating motility (Carter et al., 2008; Gibbons et al., 2005). Excess dynein-SRS_85:82_ was removed by washing the chamber twice with 15 μL MB supplemented with 1 mg/ml casein and 10 μM taxol. Fluorescently labeled microtubules were flowed and incubated in the chamber for 5 minutes and the chamber was then washed with 15 μL MB. Equal volumes of anti-GFP-coated cargo beads and 0.5 µM GFP-tagged motor proteins were mixed and incubated for 5-10 minutes on ice. 10 μL of MB with 2 mM ATP was added to the bead-motor mixture and flowed into the chamber. The cargo beads moved unidirectionally along the microtubule long axis, demonstrating that the microtubule bridges were formed by a single microtubule.

#### Data collection

Experiments were performed with a custom-built optical trapping microscope equipped with a Nikon TiE microscope body, Nikon 100× 1.49 NA plan apochromat objective, and Orca Flash 2.0 CMOS camera (Hamamatsu, Japan). LD655-labeled microtubules were excited with a 632 nm laser beam (Melles Griot) in epi-fluorescence mode. Fluorescence signal was detected by the camera with an effective pixel size of 43.3 nm after magnification. Movies were recorded at 3 Hz. The coverslip surface was scanned to identify a microtubule bridge that was stable and oriented parallel to the imaging plane between two 2-μm-diameter beads. The fluorescence image of the microtubule was brought into focus to capture the full 3D motion of the bead around the circumference of the microtubule. A 0.51-μm-diameter cargo bead freely diffusing in solution was trapped by a focused 1064 nm beam (IPG Photonics) and brought close to the microtubule. When the cargo bead exhibited unidirectional motility on the microtubule, the trapping laser was turned off, and the cargo bead was released to determine the polarity of the microtubule bridge. The cargo bead was then placed at the minus end of the microtubule to track its plus-end-directed motility throughout the entire length of the bridge. Bead movement was recorded using brightfield microscopy.

#### Analysis of bead trajectories

Center positions of the cargo beads were determined using two-dimensional Gaussian fitting in ImageJ. Z-positions of the beads were calibrated by decorating the coverslip surface with 0.51-μm-diameter beads and moving the microscope objective ±255 nm in the z direction using a PIFOC objective scanner (Physik Instrumente, Germany) with 30 nm increments. Intensities were recorded at each z-position for all selected beads to obtain a calibration curve (Fig. S2 A). Z-positions of the cargo beads in motility assays were then calculated from the calibration curve based on the intensities of the cargo beads. The 3D trajectory of each cargo bead was plotted using custom-written software in MATLAB (MathWorks) and the helical pitch and handedness of each trajectory were calculated manually.

### Quantification of spindle helicity

Spindles in early- to mid-anaphase, with two clearly separated chromosome masses but before the onset of furrowing, were chosen for analysis. Spindles were rotated so that the pole-to-pole axis was horizontal and cropped using the rectangle tool in FIJI (Schindelin et al., 2012). The positions of the two poles were manually assigned based on tubulin intensity. Spindles were resliced along the pole-to-pole axis by permuting the [x,y,z] coordinates to [y,z,x] in MATLAB. Confocal, but not light sheet, images were pre-processed in Python by subtracting an image blurred with a Gaussian kernel of standard deviation 30 pixels (scipy.ndimage.gaussian_filter1d), followed by despeckling with a 3-pixel median filter (scipy.ndimage.median_filter). Helicity was quantified using a previously published optical flow method (https://gitlab.com/IBarisic/detecting-microtubules-helicity-in-microscopic-3d-images) (Trupinic et al., 2022). Briefly, Farnebäck optical flow (Farnebäck, 2003) was calculated between each pair of successive frames lying between 30% and 70% of the pole-to-pole axis. Flow vectors were converted to polar coordinates, weighted by pixel intensities in the processed images using the “All pixels weighted helicities” method, and averaged for each spindle.

To compare helicity quantification methods in Fig. S1, the bundle tracing method was performed as previously described (Neahring et al., 2021) (a modified version of the method first published by Novak et al., 2018). For the bundle angle method, the lowest and highest plane of a z-stack were identified that contained clearly visible microtubule bundles. In these two planes, the angles of three bundles were manually measured using the line tool in FIJI and averaged. Helicity was calculated using the formula:

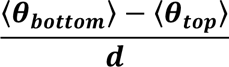

where d is the diameter of the spindle in µm.

### Quantification of spindle shape and cortical fluorescence intensity

Spindle length was determined by calculating the distance between the manually-assigned pole positions used for helicity analysis. Width was manually measured at the spindle equator from maximum intensity projections of the tubulin channel. Aspect ratio was calculated as spindle length divided by spindle width.

NuMA fluorescence intensity at the cell cortex was calculated from the single z-plane where cortical intensity appeared brightest near each spindle pole. In FIJI, short line segments 20 px (1.16 µm) wide were drawn across the cortex, through the cytoplasm, and outside the cell (background) near each spindle pole, totalling 6 line segments per cell. Using the “plot profile” function in FIJI, the maximum intensity was extracted from each line segment. The corresponding values from each pole were averaged, the background value was subtracted from the cortex and cytoplasm values, and the cortex/cytoplasm intensity ratio was calculated.

### Statistical analysis

Parametric tests were used based on the assumptions that spindle helicity, shape, and microtubule intensity ratios are approximately normally distributed, with approximately equal variance between experimental conditions. In Fig. 1 B, distributions were assessed for a significant difference from 0 helicity with one-sample t-tests using the ttest function in MATLAB. Helicities, cortical intensities, and spindle shape parameters between experimental conditions were compared using one-way ANOVA with post-hoc Tukey-Kramer tests, using the anova1 and multcompare functions in MATLAB, or using two-sample t-tests with the ttest2 function in MATLAB when only two groups were compared. We used p < 0.05 as a threshold for statistical significance, and all tests were two-tailed. Number of cells, number of independent experiments, and p-values are provided in figure legends.

## Acknowledgements

We thank Ivan Ivanov, Talon Chandler, and Shalin Mehta for experimental assistance, and Stefan Diez, Laura Meißner, Iain Cheeseman, Stefan Grill, Orion Weiner, Arthur Molines, and members of the Yildiz and Dumont labs for helpful discussions. This work was supported by NIH (R35GM136420, S.D.; R35GM094522, A.Y.; F31CA275394, N.H.C.), NSF (1554139, S.D.; MCB-1617028 and MCB-1055017, A.Y.), NSF 1548297 Center for Cellular Construction, the Chan Zuckerberg Biohub, the UCSF Byers Award, and the UCSF Program in Breakthrough Biomedical Research (S.D.); NSF Graduate Research Fellowships (L.N. and C.R.); the Fannie and John Hertz Foundation Fellowship (L.N.); the American Heart Association Predoctoral Fellowship (N.H.C.); and the UCSF Discovery Fellows Program (L.N., N.H.C., C.R.).

## Author contributions

Conceptualization, L.N., A.Y., S.D.; data curation, L.N., Y.H.; formal analysis, L.N., Y.H., G.L.; funding acquisition, S.U., A.Y., S.D.; investigation, L.N., Y.H., N.H.C., G.L., C.J.R.; methodology, L.N., Y.H., N.H.C., G.L.; resources, L.N., G.L., J.F., K.N., R.S., S.U.; software, L.N., G.L.; supervision, L.N., R.S., S.U., A.Y., S.D.; validation, L.N., Y.H., N.H.C., G.L.; visualization, L.N., Y.H., G.L., S.U.; writing – original draft, L.N., Y.H.; writing – review & editing, L.N., Y.H., N.H.C., G.L., J.F., C.J.R., K.N., R.S., S.U., A.Y., S.D.

## Figure legends

**Figure S1. Comparison of helicity quantification methods and comparison of helicity with spindle shape. (A)** Schematic diagrams illustrating three methods of quantifying spindle helicity. The optical flow method is calculated from all pixels in an end-on (XZ) view of the spindle, the bundle tracing method is calculated from ∼10 bundles per cell manually traced from an end-on (XZ) view of the spindle, and the bundle angle method is calculated from 6 bundles in a top-down (XY) view of the spindle. **(B)** Scatterplots comparing the same 18 siCtrl (gray) and 14 siLGN (black) cells analyzed by each of the three quantification methods. Pearson’s correlation coefficients are shown on each plot. **(C-E)** Violin plots with internal boxplots of spindle length (C), width (D), and aspect ratio (length/width, E) for all spindles analyzed in all conditions. n.s. not significant, *p<0.05, **p<0.005, ***p<0.0005, ****p<0.00005, one-way ANOVA with Tukey’s post-hoc test. **(F-H)** Scatterplots of spindle length (F), width (G), and aspect ratio (H) vs. spindle helicity, averaged for each experimental condition. Vertical and horizontal gray lines indicate standard deviation. Helicity is not significantly correlated with spindle length, width, or aspect ratio (two-sided Pearson’s correlation test). Pearson’s correlation coefficients are provided in the lower right of each plot.

**Figure S2. Validation of midzone kinesin experiments. (A)** in the suspended microtubule bridge assay, the z-position of the cargo bead was calibrated by measuring the average intensity of 17 surface-immobilized cargo beads by scanning the microscope objective from -255 nm to 255 nm relative to the imaging plane. Error bars represent SD. The intensity profile was fitted to a third-order polynomial (red curve, R^2^ = 0.9995) to obtain the calibration curve. **(B-D)** Western blots of KIF4A (B), MKLP1 (C), or KIF4A and MKLP1 (D) levels in MCF10A cells transfected with siRNA targeting luciferase (control), KIF4A, and/or MKLP1 for 48 hours, as indicated. GAPDH is shown as a loading control. **(E-F)** Live confocal images of the late anaphase (E) and telophase (F) phenotypes after KIF4A or MKLP1 knockdown. In late anaphase, siKIF4A spindles over-elongate and have poorly organized midzone bundles. In telophase, the midbody is extended (siKIF4A) and cells have cytokinesis defects (siMKLP1). Scale bars = 5 µm. Brightfield (upper row) and SiR-tubulin (lower row) images represent single z-planes **(G)** Helicity of anaphase spindles in MCF10A cells treated with 0.1% DMSO or 10 µM STLC, calculated from SiR-tubulin intensity. Black lines represent mean ± SD. n = 42 STLC-treated spindles pooled from N = 4 independent experiments, and the same 54 DMSO-treated spindles shown in Fig. 3 B. n.s., not significant, two-sample t-test.

**Figure S3. Characterization of dynein and LGN depletions. (A)** Western blot of dynein intermediate chain levels in MCF10A cells transfected with siRNA targeting luciferase (siCtrl) or dynein heavy chain for 48 hours. Depletion of dynein intermediate chain is correlated with dynein heavy chain depletion (Levy and Holzbaur, 2008). GAPDH is shown as a loading control. **(B)** Western blot of LGN in MCF10A cells transfected with siRNA targeting luciferase (siCtrl) or LGN for 48 hours. GAPDH is shown as a loading control. **(C)** Immunofluorescence images (single z-planes) of anaphase MCF10A cells stained for NuMA (magenta) and DNA (blue). Scale bars = 5 µm. **(D)** Quantification of NuMA enrichment at the cell cortex relative to NuMA intensity in the cytoplasm (see Materials and methods), in immunofluorescence images of MCF10A cells transfected with siRNA targeting luciferase (control) or LGN. Dashed line indicates a cortex/cytoplasm ratio of 1, i.e. no cortical enrichment. Black lines represent mean ± SD. n = 22 cells from 2 independent days in each condition. ****p = 1.55×10^-10^, two-sample t-test.

