## Supplemental Figures for "Torques within and outside the human spindle balance twist at anaphase"

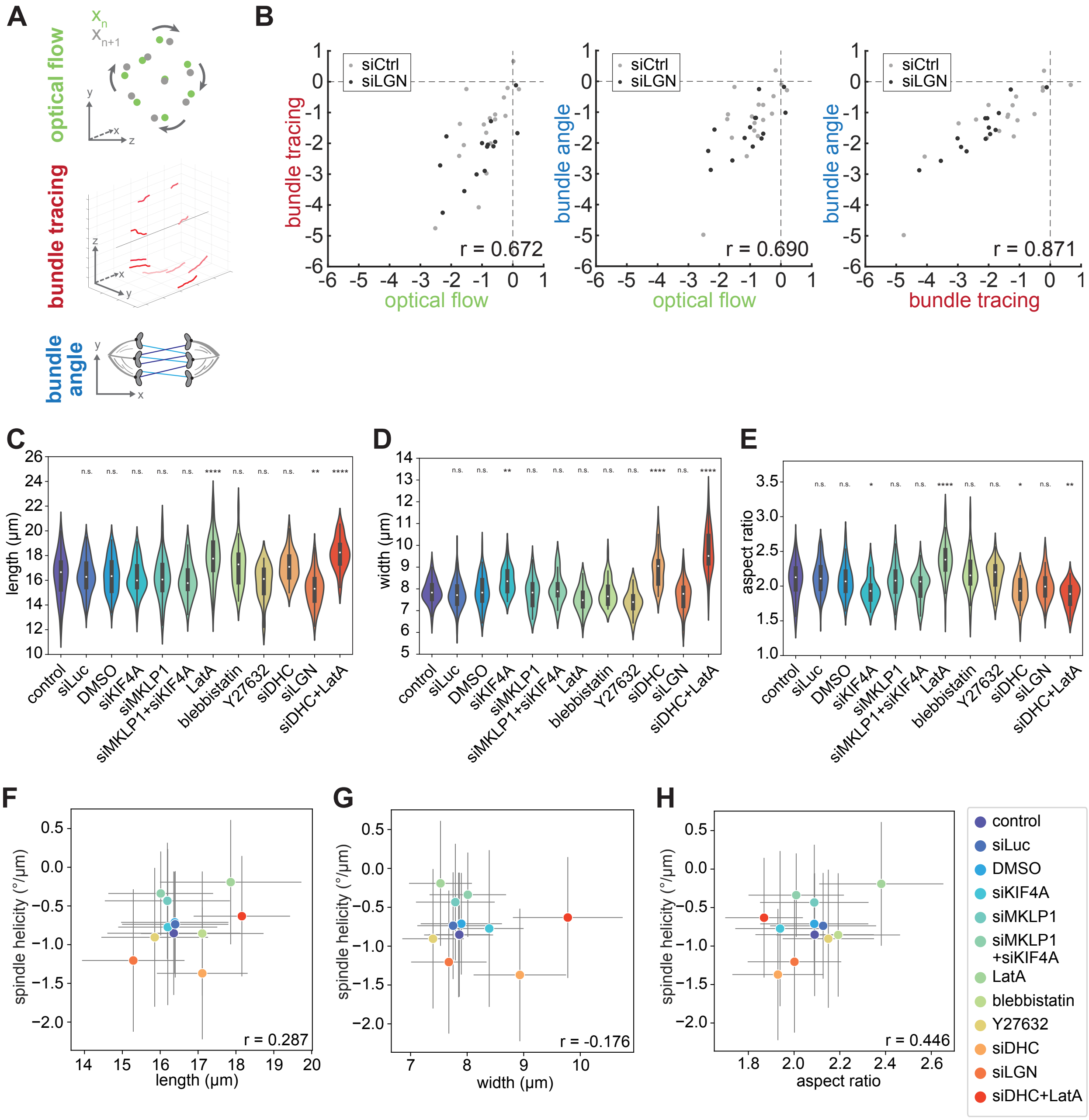


**Figure S1. Comparison of helicity quantification methods and comparison of helicity with spindle shape**

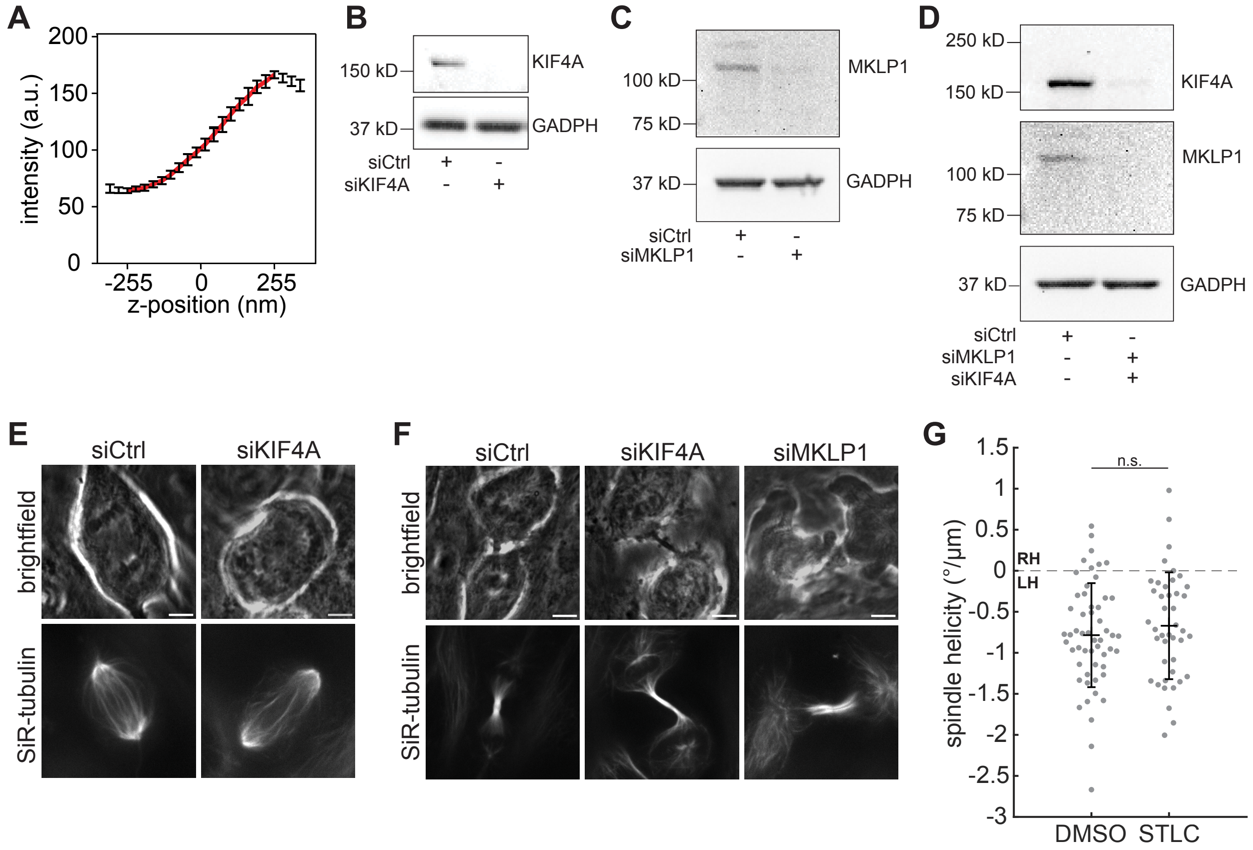


**Figure S2. Validation of midzone kinesin experiments**

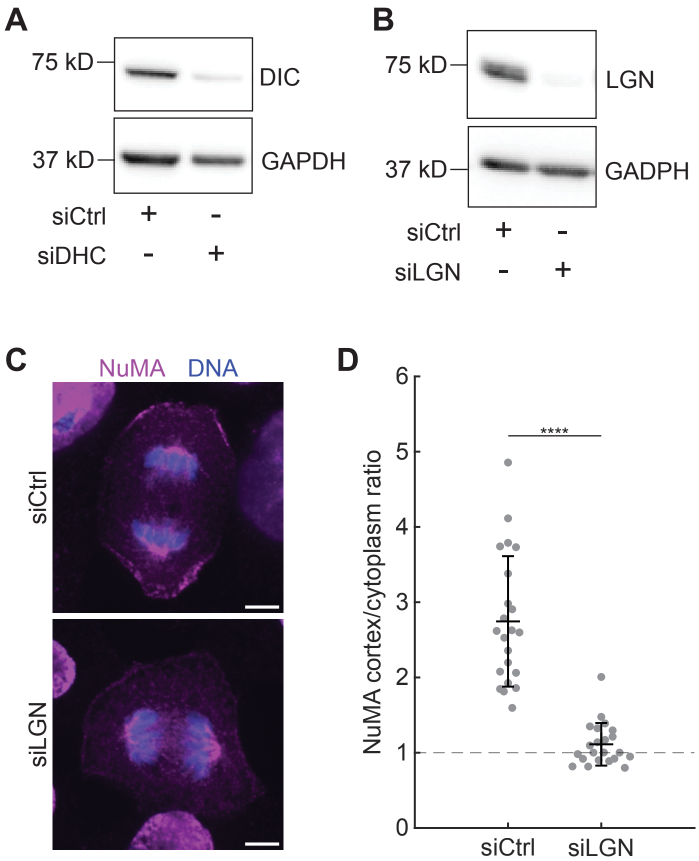


**Figure S3. Characterization of dynein and LGN depletions**
